# Two-step binding kinetics of tRNA^Gly^ by the *glyQS* T-box riboswitch and its regulation by T-box structural elements

**DOI:** 10.1101/350413

**Authors:** Jiacheng Zhang, Bhaskar Chetnani, Eric D. Cormack, Wei Liu, Alfonso Mondragón, Jingyi Fei

## Abstract

T-box riboswitches are *cis*-regulatory RNA elements that regulate mRNAs encoding for aminoacyl tRNA synthetases or proteins involved in amino acid biosynthesis and transport. Rather than using small molecules as their ligands, as do most riboswitches, T-box riboswitches uniquely bind tRNA and sense their aminoacylated state. Whereas the anticodon and elbow regions of the tRNA interact with Stem I, located in the 5’ portion of the T-box, sensing of the aminoacylation state involves direct binding of the NCCA sequence at the tRNA 3’ end to the anti-terminator sequence located in the 3’ portion of the T-box. However, the kinetic trajectory that describes how each of these interactions are established temporally during tRNA binding remains unclear. Using singlemolecule fluorescence resonance energy transfer (smFRET), we demonstrate that tRNA binds to the riboswitch in a two-step process, first with anticodon recognition followed by NCCA binding, with the second step accompanied by an inward motion of the 3’ portion of the T-box riboswitch relative to Stem I. By using site-specific mutants, we further show that the T-loop region of the T-box significantly contributes to the first binding step, and that the K-turn region of the T-box influences both binding steps, but with a more dramatic effect on the second binding step. Our results set up a kinetic framework describing tRNA binding by T-box riboswitches and highlight the important roles of several T-box structural elements in regulating each binding step.

**SIGNIFICANCE:** Bacteria commonly use riboswitches, *cis*-regulatory RNA elements, to regulate the transcription or translation of the mRNAs upon sensing signals. Unlike small molecule binding riboswitches, T-box riboswitches bind tRNA and sense their aminoacylated state. T-box modular structural elements that recognize different parts of a tRNA have been identified, however, how each of these interactions is established temporally during tRNA binding remains unclear. Our study reveals that tRNA binds to the riboswitch in a two-step mechanism, with anticodon recognition first, followed by binding to the NCCA sequence at the 3’ end of the tRNA with concomitant conformational changes in the T-box. Our results also highlight the importance of the modular structural elements of the T-box in each of the binding steps.

## INTRODUCTION

Riboswitches are *cis*-regulatory RNA elements that recognize and respond to defined external signals to affect transcription or translation of downstream messenger RNAs (mRNAs) (1–3). Riboswitches generally consist of two domains: a sensory or aptamer domain and a regulatory domain or expression platform. The expression platform can adopt different conformations in response to ligand binding to the aptamer, and in this way controlling gene expression outcome (1). The aptamer of each riboswitch class contains conserved sequence motifs and unique secondary or tertiary structural elements that help distinguish and bind specific ligands (4). Bacterial T-box riboswitches represent a unique class of riboswitches that do not bind small molecule ligands, instead they recognize and bind tRNA molecules and sense directly their aminoacylation state (5). T-box riboswitches serve as excellent paradigms to understand RNA-RNA interactions and RNA-based regulation.

T-box riboswitches are found in Gram-positive bacteria and are usually located in the region upstream of mRNA sequences encoding aminoacyl tRNA synthetases and proteins involved in amino acid biosynthesis and transport and hence participate directly in amino acid homeostasis (5). In general, the aptamer domain of all T-box riboswitches contains a long stem, Stem I, responsible for specific tRNA binding (6). The expression platform can adopt either a terminator or anti-terminator conformation, depending on whether the bound tRNA is charged or uncharged (5, 7). In most T-box riboswitches, binding of a charged tRNA to the T-box leads to rho independent transcription termination whereas an uncharged tRNA stabilizes the antiterminator conformation and leads to transcription read-through (5, 7). Whereas Stem I and the anti-terminator domain are highly conserved among T-box riboswitches, the region connecting them can vary. The *Bacillus subtilis glyQS* T-box riboswitch, involved in glycine regulation, represents one of the simplest T-box riboswitches (8). Only a short linker and a small stem, Stem III, connect Stem I and the anti-terminator domain (Fig. 1).

**Figure 1.**
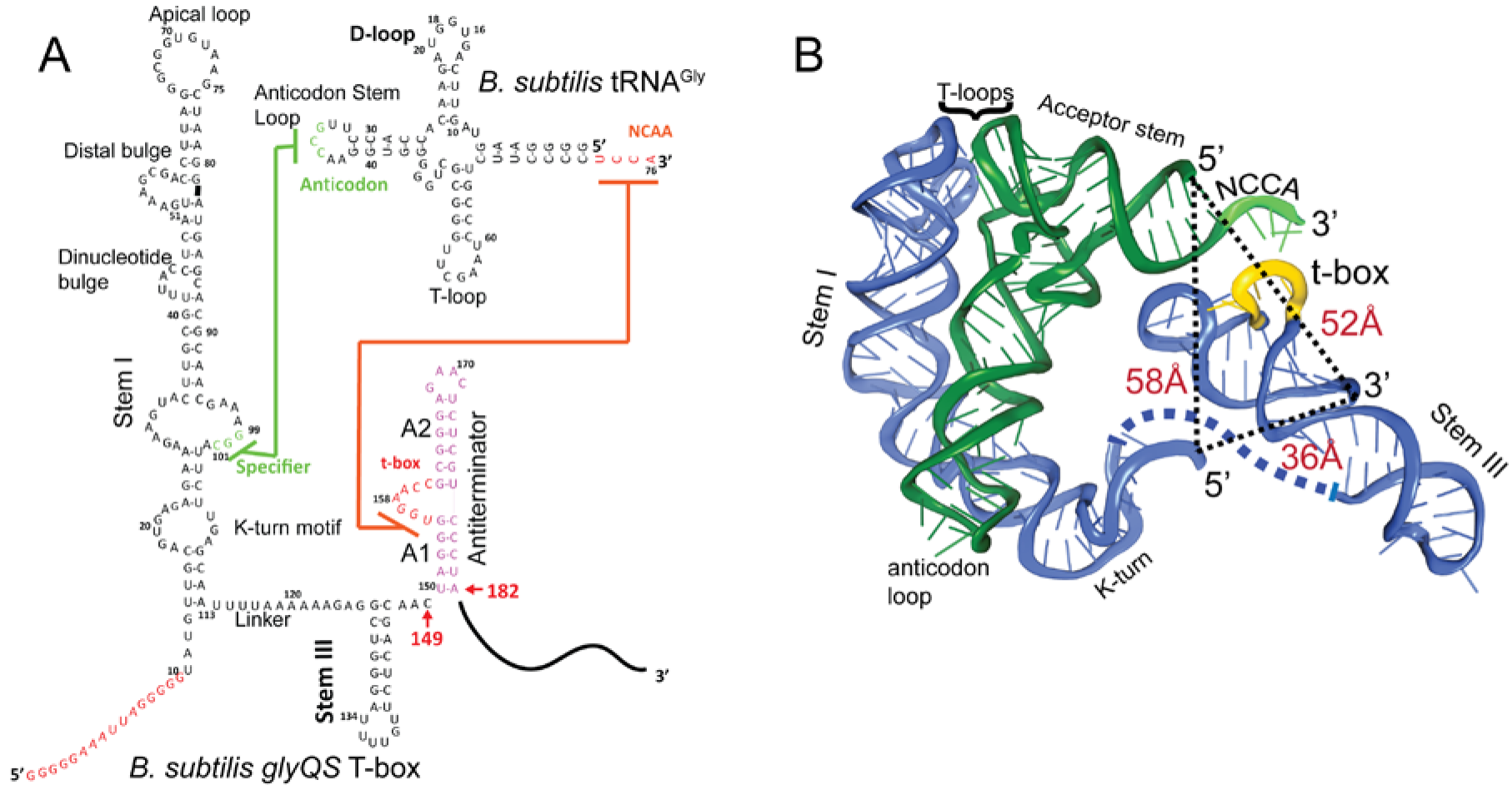
Secondary and tertiary structures of *B. subtilis glyQS* T-box riboswitch and tRNA^Gly^. **(A)** Secondary structure diagrams of the *B. subtilis glyQS* T-box riboswitch and *B. subtilis* tRNA^Gly^ used in this study. Green and orange lines indicate interactions between the T-box specifier loop and the tRNA anticodon and between the T-box t-box sequence and the tRNA 3’ NCCA, respectively. The sequence of *glyQS* T-BOX in red is added for surface immobilization. **(B)** Ribbon diagram of a model of a complex between the *B. subtilis glyQS* T-box riboswitch (blue) and *B. subtilis* tRNA^Gly^ (green) based on SAXS data (17). Distances between the 5’ and 3’ ends of the T-box and the 5’ end of the tRNA^Gly^ are shown (black dash lines). The NCCA sequence at the 3’ end of the tRNA is shown in light green and the t-box sequence in the T-box is shown in yellow.

Recognition of tRNA by a T-box riboswitch involves three main structural elements of the tRNA: the anticodon region, the “elbow” region formed by the conserved T-and D-loops, and the 3’ NCCA sequence (Fig. 1B). The anticodon and elbow regions of the tRNA interact with Stem I directly. Stem I contains several phylogenetically conserved structural motifs (6), including a K-turn motif, a specifier loop, a distal bulge, and an apical loop (6) (Fig. 1). Bioinformatics and structural analyses have collectively revealed the interactions between Stem I and the tRNA (9, 11). Specifically, the co-crystal structures of Stem I/tRNA complexes show that Stem I flexes to follow closely the tRNA anticodon stem and interacts directly with the anticodon loop and the elbow through its proximal and distal ends, respectively (11). The distal bulge and the apical loop fold into a compact structural module of interdigitated T-loops (12, 13), which interact directly with conserved unstacked nucleobases at the tRNA elbow (9, 11). In addition, the structures revealed that Stem I turns sharply around two hinge regions using a conserved dinucleotide bulge and the K-turn motif (11, 14). Sensing of the aminoacylation state involves direct binding of the tRNA 3’ end to a highly conserved bulge in the T-box, the t-box sequence (15) (Fig. 1). A free NCAA end can base pair with the t-box sequence, enabling the anti-terminator conformation, whereas a charged NCCA end prevents the formation of the t-box/NCCA interactions, leading to the more stable terminator conformation (5, 7). Importantly, discrimination between the charged and uncharged tRNA does not require any additional proteins, such as EF-Tu (16), and is driven solely by RNA/RNA interactions.

Although there are no atomic-level structural details on the interactions between tRNA and the anti-terminator region, Small Angle X-ray Scattering (SAXS)-derived models of the entire *B. subtilis glyQS* T-box riboswitch in complex with tRNA are available (17, 18). The two models are distinct, one presenting a more compact structure where all the previously observed interactions between Stem I and tRNA are preserved and the 3’ NCAA sequence of the tRNA helps stabilize a coaxial stem formed by Stem III and the anti-terminator region (17), while the second model shows a more extended and relaxed structure where the interactions with the anticodon are preserved but the contacts with the tRNA elbow are not present (18). In addition, there is a dearth of information on the kinetics of the binding process. Whereas it is clear that tRNA recognition involves several specific interactions, their binding temporal sequence remains elusive. In addition, it is unclear whether sensing of the 3’ end of the tRNA involves any additional conformational changes in the T-box. Here, by introducing donor-acceptor fluorophore pairs at several locations in the tRNA and the T-box riboswitch, and using single-molecule fluorescence resonance energy transfer (smFRET), we demonstrate the temporal order of events in the trajectory of tRNA binding. Our results demonstrate that tRNA binds to the riboswitch in two steps, with its anticodon being recognized first, followed by NCCA binding accompanied with an inward motion of the 3’ region of the T-box riboswitch, including Stem III and the anti-terminator stem, relative to Stem I. In addition, by introducing mutations at different locations of the T-box, we further show that the two-step binding kinetics is regulated by structural elements in the T-box riboswitch.

## RESULTS

### Binding of cognate tRNA by the glyQS T-box results in two distinct FRET states

To observe directly the binding of tRNA to the T-box, we placed the donor dye (Cy3) on the 3’ end of a T-box fragment (T-box_182_), and the acceptor dye (Cy5) on the 5’ end of the tRNA^Gly^, where the subscript “182” denotes the length of the T-box construct (*SI Appendix,* Figs. 1A and S1). T-box_182_ spans Stem I, the linker sequence, Stem III and the anti-terminator, but does not contain the terminator sequence, thereby preventing the transition to the terminator conformation. A short RNA extension sequence was added to the 5’ end of the T-box for surface immobilization (Fig. 2A and *SI Appendix,* Fig. S1). Single-molecule fluorescence images were recorded under equilibrium condition in the presence of 30nM tRNA^Gly^-Cy5. Binding of tRNA^Gly^-Cy5 results in a major distribution of FRET values around 0.7, with 79±4% of the traces showing a stable signal at 0.7 and (9±5)% traces sampling from 0.7 to 0.4 (Fig. 2B and C). The SAXS model (17) predicts a distance between the labeling positions at the 3’ end of the T-box_182_ and the 5’ end of the tRNA to be around 52 Å. (Fig. 1B). Based on a Förster distance of 54-60 Å (19, 20), our measured FRET value is within the range of estimated FRET values (0.56-0.7). Therefore, we assign the 0.7 FRET state to be the fully bound state of the tRNA^Gly^ by the T-box.

**Figure 2.**
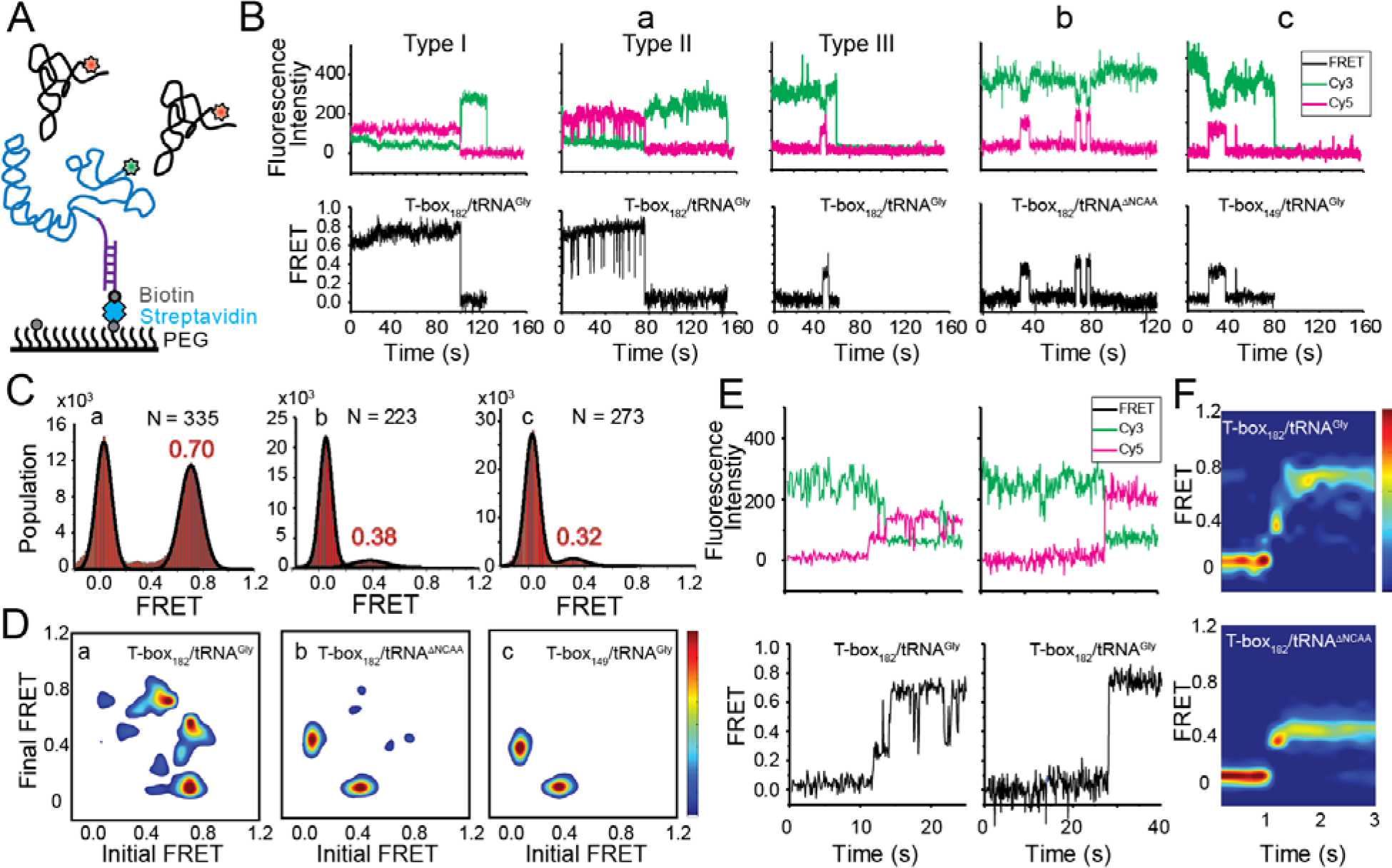
Two-step binding of uncharged tRNA to the *glyQS* T-box riboswitch. **(A)** FRET labeling scheme for the T-box and tRNA. Cy3 (green star) and Cy5 (red star) fluorophores are attached at the 3’ of the T-box (blue) and the 5’ of the tRNA (black), respectively. *glyQS* T-box riboswitch molecules are anchored on slides through a biotinylated DNA probe (purple) hybridized to a 5’ extension sequence on the T-box. **(B)** smFRET *vs.* time trajectories of T-box_182_-Cy3(3’) with (a) tRNA^Gly^-Cy5 or (b) tRNA^ΔNCCA^-Cy5 and (c) T-box_149_-Cy3(3’) with tRNA^Gly^-Cy5. Cy3 and Cy5 fluorescence intensity traces (upper panel), and their corresponding smFRET traces calculated as I_oy5_ / (I_oy3+_I_Cy5_) (lower panel). **C)** One-dimensional FRET histograms of combinations (a), (b) and (c) as described above. FRET peaks are fit with a Gaussian distribution (black curve) and the peak centers are shown in red. “N” denotes the total number of traces in each histogram. **D)** Transition density plot (TDP) of combinations (a), (b) and (c). Contours are plotted from blue (lowest population) to red (highest population). **E)** Representative smFRET trajectories showing real-time binding of tRNA^Gly^-Cy5 to T-box_182_-Cy3(3’) in a steady-state measurement. Traces showing transitions from unbound state (0 FRET) to partially bound state (0.4 FRET) to fully bound state (0.7 FRET) (left) and unbound state directly to fully bound state (right). **F)** Surface contour plot of time-evolved FRET histogram of T-box_182_-Cy3(3’) with tRNA^Gly^-Cy5 (top) and tRNA^ΔNCCA^-Cy5 (bottom). Traces that reach the 0.7 FRET state (cutoff >0.55) are included in the plot for tRNA^Gly^-Cy5 to reveal better the transition from the 0.4 to the 0.7 FRET state. Time-evolved FRET histograms of all traces are shown in *SI Appendix*, Fig. S5D for comparison.

In order to assign the 0.4 FRET value to specific tRNA binding states, tRNA^Tyr^-Cy5 and tRNA^ΔNCCA^-Cy5 (“ΔNCCA” denotes a tRNA^Gly^ with deleted 3’ NCCA sequence) were flowed in the flow-chamber with pre-immobilized T-box_182_-Cy3(3’) (3’ denotes that the label was added at the 3’ end). We did not observe any binding of tRNA^Tyr^-Cy5 (*SI Appendix*, Fig. S2), confirming that recognition of the anticodon by the specifier region is required for tRNA binding. In the presence of tRNA^ΔNCCA^-Cy5, we observed a fluctuating signal between 0.4 and 0 FRET (Fig. 2B and D), with an average lifetime of the 0.4 FRET state of 3.6±0.6 s and an average waiting time before binding of 31.3±5.3 s (*SI Appendix*, Fig. S3B). Taken together with the results from the tRNA^Gly^, tRNA^ANCCA^ and tRNA^Tyr^ binding experiments, we assign the 0.4 FRET state to a partially bound state where only the anticodon interactions have been established.

To further confirm the assignment of the FRET states, we generated T-box_149_, where the antiterminator sequence is truncated (Figs. 1A and *SI Appendix*, S1). Based on the structure model from the SAXS data (17) we predicted that a Cy3 dye placed either at the end of Stem III (T-box^149^) or at the end of the anti-terminator stem (T-box_182_) are localized in close proximity in three dimensions, further confirmed by the distance measurement using smFRET (*SI Appendix*, Fig.S4). Therefore, we expect that if tRNA^Gly^-Cy5 can reach the same fully bound state in T-box_149_ as in T-box_182_, a high FRET state centered at 0.7 would be observed. However, using T-box_149_-Cy3(3’) in combination with tRNA^Gly^-Cy5, we again observed transient binding of tRNA^Gly^ with a FRET value centered at ~0.4 with the same average lifetime as observed with the T-box_182_-Cy3(3’) and tRNA^ΔNCCA^-Cy5 combination (Fig. 2B-D and *SI Appendix*, Fig. S3B and C). Therefore these two complexes (T-box_182_ + tRNA^ΔNCCA^ and T-box_149_ + tRNA^Gly^) represent the same binding state of the tRNA, i.e. the state where binding of the anticodon to the specifier region has been established, but is unstable without the further interactions between the NCCA and the t-box region.

Collectively, our results suggest a two-step binding model where the establishment of the interaction with the anticodon precedes the interactions with the NCCA. Without the interaction between the NCCA and the t-box sequence the binding of tRNA^Gly^ is not stable. From the binding kinetics of tRNA^ΔNCCA^, we estimated the association rate constant (*k*_1_) and the disassociation rate constant (*k*_−1_) for the first binding step to be (5.0±1.7)X10^5^ s^−1^·M^−1^ and 0.28±0.04 s^−1^, respectively (Figs. 5 and 6E and *SI Appendix*).

**Figure 5.**
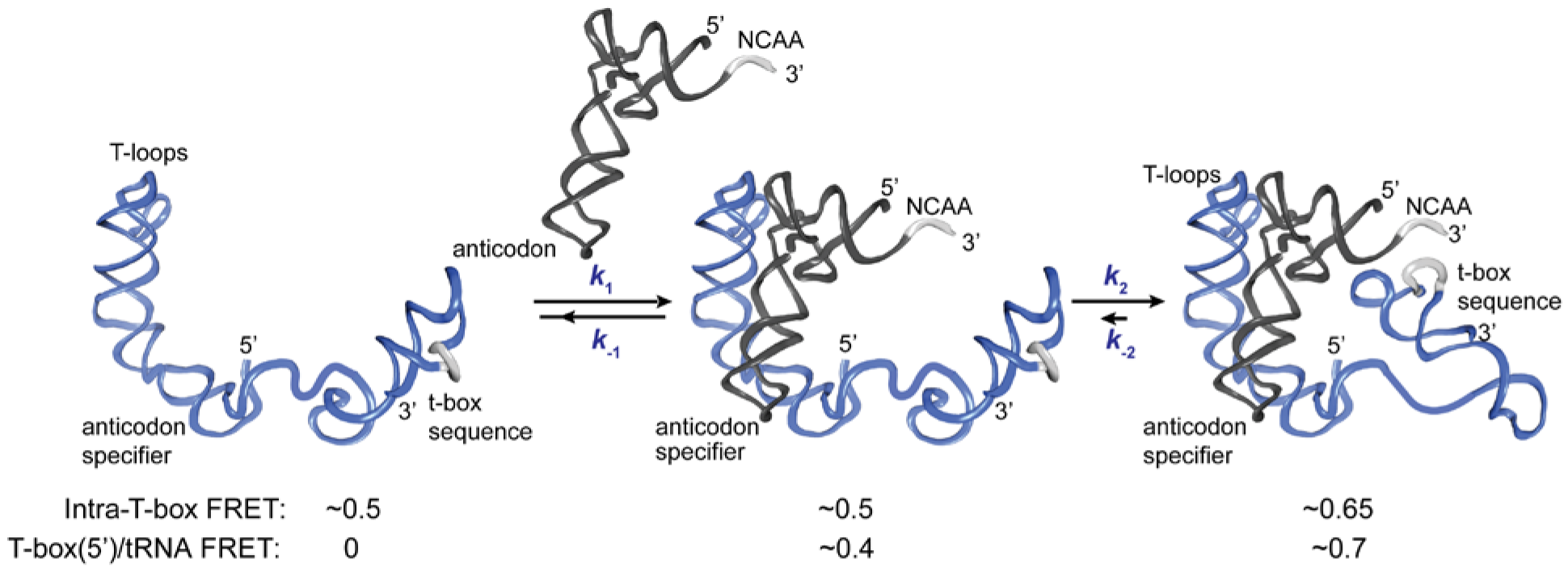
Kinetic model for the two-step binding of *glyQS* T-box riboswitch and uncharged tRNA^Gly^. Details of the model are described in the text. Rate constants are summarized in Fig. 6E.

### The transition from anticodon recognition to NCCA binding is rapid for uncharged tRNA

We classified smFRET traces for T-box_182_-Cy3(3’) in complex with tRNA^Gly^-Cy5 into three types (Fig. 2B): (I) traces stably sampling the 0.7 state (79±4%), (II) traces transiently transitioning from the 0.7 state to the 0.4 state (9±5%), and (III) traces only sampling the 0.4 state without reaching 0.7 state (12±5%). The low percentage of Type III traces indicates that once the anticodon is recognized, the commitment to the next binding step, NCCA interactions, is high. The majority of the traces showed that the tRNA^Gly^ remained mostly in the fully bound state (Type I) until the fluorophore photobleached, with the actual lifetime limited by photobleaching (T_0.7_ > 24 s, where T_0.7_ denotes the lifetime of the 0.7 FRET state) (*SI Appendix*, Fig. S3A). The observation that tRNA^Gly^ is able to transit from the fully bound state back to the partially bound state (Type II) suggests that the NCCA/t-box interaction can break occasionally (Fig. 2B). We estimated the lifetime of the transiently sampled partially bound state to be 0.35±0.09 s (*SI Appendix*, Fig. S3A), ~10-fold shorter than the partially bound state without the NCCA end.

While the majority of the T-box molecules were already bound to tRNA^Gly^ before starting data acquisition, we could detect that some molecules show real-time binding during imaging acquisition. We observed only a few traces briefly sampling the 0.4 FRET state from the zero FRET (unbound) state before reaching the 0.7 FRET state, while most traces directly sampled the 0.7 FRET state without a detectable 0.4 FRET, likely due to our imaging time resolution (100 ms per frame). We post-synchronized the FRET traces at the transition point from the zero FRET state to the first sampled 0.4 FRET state, and plotted in a time-evolved FRET histogram. From the time-evolved FRET histogram (Fig. 2F), we estimated roughly that the upper limit of the lifetime spent at the 0.4 FRET state is ~100 ms, very rapidly followed by establishment of NCCA/t-box interactions. In contrast, tRNA^ΔNCCA^ could not pass the 0.4 FRET state. To capture better realtime binding, we performed a flow experiment, where tRNA^Gly^-Cy5 was flowed in to a chamber with immobilized T-box_182_-Cy3(3’) during imaging acquisition. The corresponding post-synchronized time-evolved FRET histogram again shows a fast transition into the fully bound state (*SI Appendix*, Fig. S5). In addition, the association rate constant of tRNA^Gly^ in the real-time flow experiment is (7.5±0.7)×10^5^ s^−1^·M^−1^, consistent with the *k*_1_ of tRNA^ΔNCCA^ and confirming that the NCCA end of the tRNA does not participate in the first binding step.

**Figure 4.**
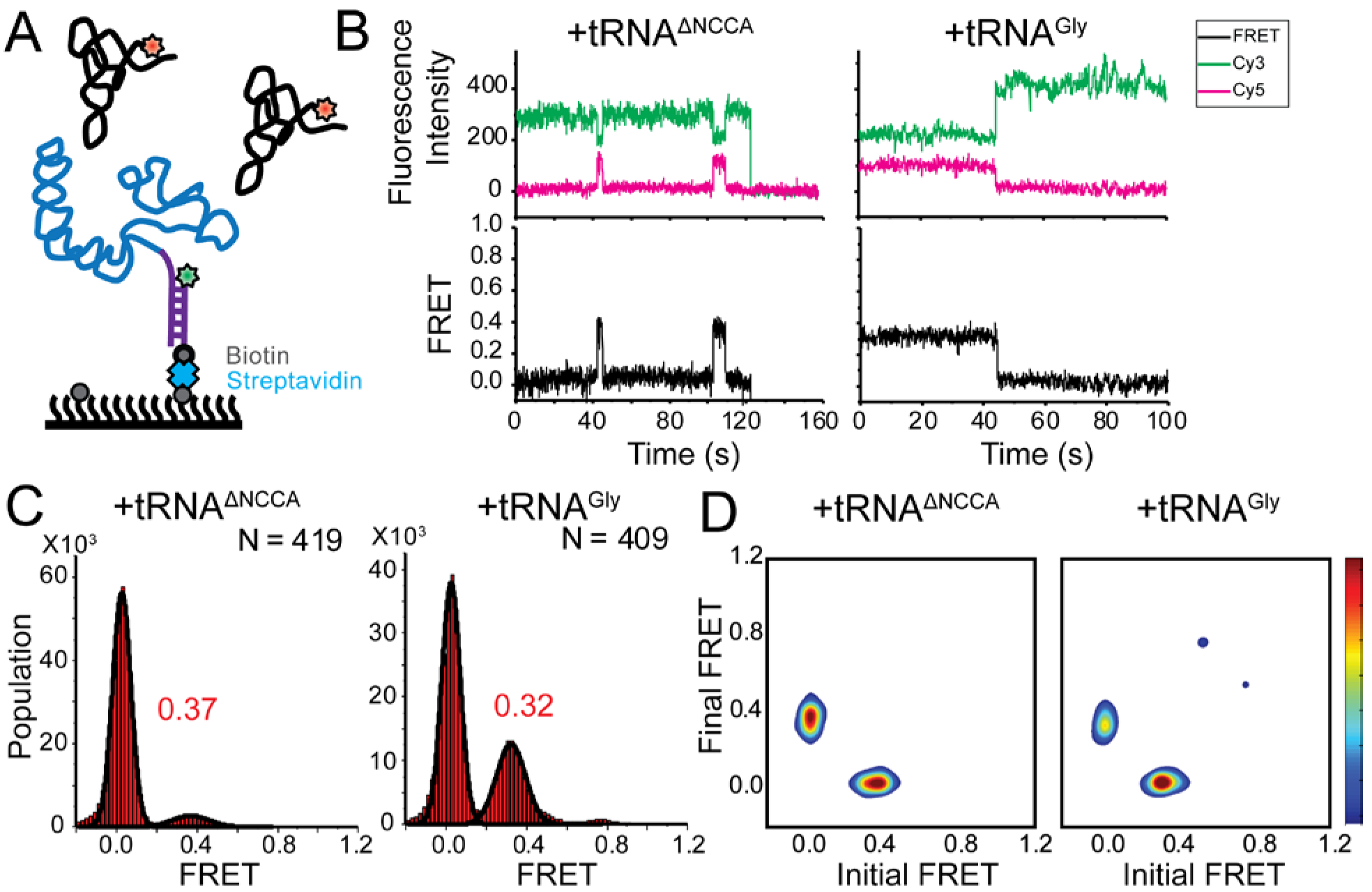
FRET between fluorophores at the 5’ end of the *glyQS* T-box riboswitch and 5’ end of tRNA^Gly^ is insensitive to the two binding states. **(A)** Cy3 (green star) and Cy5 (red star) are attached at the 5’ extension of the T-box (blue) and the 5’ of the tRNA (black), respectively. **(B)** smFRET trajectories of T-box-Cy3(5’) with tRNA^ΔNCCA^-Cy5 (left) and tRNA^Gly^-Cy5 (right). **(C)** One-dimensional FRET histograms of T-box-Cy3(5’) with tRNA^ANCCA^-Cy5 (left) and tRNA^Gly^-Cy5 (right). **(D)** TDP of T-box-Cy3(5’) with tRNA^ΔNCCA^-Cy5 (left) and tRNA^Gly^-Cy5 (right)

From the real-time binding kinetics of tRNA^Gly^ to T-box_182_, we estimated a transition rate constant from the partially bound state to the fully bound state (*k*_2_) of ~10 s^−1^ (Fig. 2F and *SI Appendix*, Fig. S5). On the other hand, as transitions back to the partially bound state from the fully bound state were only observed in ~10% traces, we interpreted this to mean that the reverse transition rate constant (*k*_−2_) is very small, and the second binding step in the wild-type (WT) T-box with uncharged tRNA^Gly^ is close to irreversible (Figs. 5 and 6E).

### Establishment of the NCCA/t-box interaction is accompanied by conformational changes in the T-box riboswitch

We next investigated whether tRNA binding requires any conformational changes in the T-box itself. Using doubly labeled T-box_182_, with Cy3 at the 3’ end and Cy5 at the 5’ hybridization extension, we observed a high FRET state (centered at ~0.75) in the absence of tRNA (*SI Appendix*, Fig. S4). Based on the structural model (17), we estimated the distance between the 5’ and 3’ ends of the T-box_182_ to be ~36 Å (Fig. 1B). Our measured FRET value is slightly less than the predicted FRET value (~0.9), likely due to the engineered 5’ extension sequence used to immobilize the T-box. No noticeable change was detected upon incubation with unlabeled tRNA^Gly^ (*SI Appendix*, Fig. S4C), indicating that the 3’ portion (Stem III plus the anti-terminator stem) does not move away from the 5’ portion (Stem I). Given that the measured FRET efficiency of 0.75 is already located beyond the FRET sensitive region, it is unlikely that any inward motion of the 3’ portion relative to the 5’ could be detected. To overcome this limitation, we added extensions at both the 3’ and 5’ ends (Fig. 3A and *SI Appendix*, Fig. S1). ITC experiments suggest that addition of a 5’ and/or a 3’ extension sequences to the T-BOX does not affect tRNA binding (*SI Appendix*, Fig. S5). With this intra-T-box FRET scheme, we observed a FRET shift from ~0.5 to ~0.65 when tRNA^Gly^ was added (Fig. 3B), indicating that the 3’ half of the T-box moved closer to the 5’ half, potentially with the T-box becoming more compact due to the presence of the cognate tRNA^Gly^. Adding non-cognate tRNA^Phe^ or tRNA^ΔNCCA^ gave similar FRET values as the T-box alone (Fig. 3B), suggesting that the conformational change is associated with binding of the NCCA, not with anticodon recognition.

**Figure 3.**
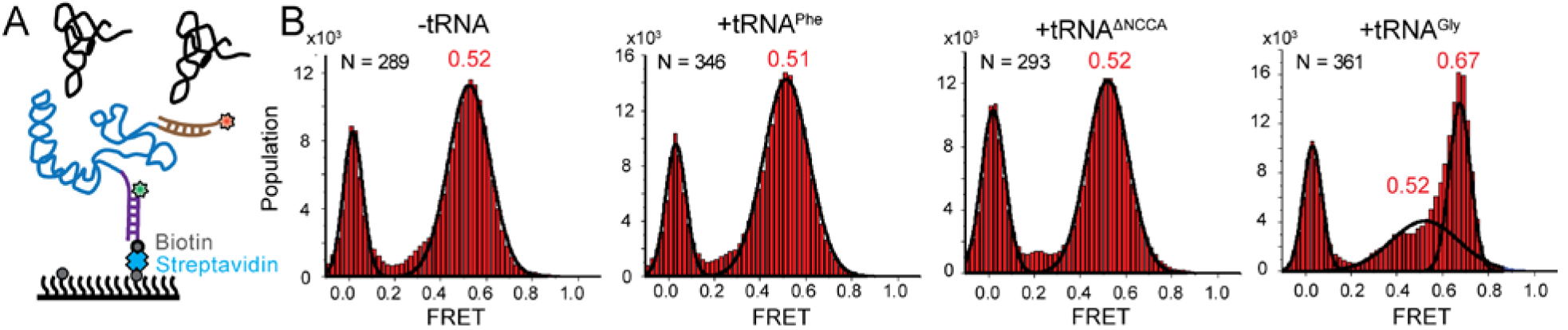
Conformational changes following tRNA binding in the *glyQS* T-box riboswitch revealed by an intra-T-box FRET pair. **(A)** Intra-T-box FRET scheme. Cy3 (green star) and Cy5 (red star) are attached at the 5’ and 3’ extensions of T-box (blue), respectively. **(B)** Onedimensional FRET histograms of T-box_182_ alone, with tRNA^Phe^, with tRNA^ΔNCCA^, and with tRNA^Gly^.

### The hinge of the intra-T-box conformational change is located near the K-turn region

Based on the known structures of Stem I in complex with tRNA^Gly^ (11, 14), it was hypothesized that the intra-T-box conformational change associated with the NCCA interaction is likely to involve the K-turn region. If this were the case, FRET between labels on the tRNA and near the K-turn region will be insensitive to the two binding states of the tRNA. Based on structural studies (11, 14, 17) (Fig. 1B), the 5’ end of the T-box is in close proximity to the K-turn. We measured FRET between a Cy3 placed at the 5’ end of the T-box (T-box_182_-Cy3(5’)) and tRNA^Gly^-Cy5 (Fig. 4A). As predicted, using this FRET pair binding of both tRNA^Gly^ and tRNA^ΔNCCA^ generated a similar FRET value centered at ~0.35 (Figs. 4B and C). The FRET traces behaved differently for these two tRNA molecules. For tRNA^ΔNCCA^, the signal fluctuated between zero and 0.35 (Fig. 4D), with a lifetime of the 0.35 FRET state of 4.5±1.0 s, reminiscent of the 0.4 FRET state using the tRNA/T-box_182_-Cy3(3’) FRET pair. For tRNA^Gly^, the signal was more stably centered at 0.35 (Fig. 4B). Since the tRNA-Cy5/T-box_182_-Cy3(5’) FRET pair cannot distinguish the partially bound from the fully bound state, we fit the lifetime with a double-exponential decay. The fast dissociation fraction has a lifetime of 5.1±0.2 s, consistent with the lifetime for the partially bound state, and the low dissociation fraction has a lifetime of 38.4±5.2 s, representing the stable fully bound state. Overall, the measurements with the tRNA-Cy5/T-box-Cy3(5’) FRET pair further validate the two-step binding model, and support the prediction that the hinge of the intra-T-box conformational change is roughly located near the K-turn region.

### A mutation in the T-loop region affects the first binding step but has minimal effect on the second binding step

The interdigitated T-loops structure formed by the interactions between the distal bulge and the apical loop at the distal end of Stem I has been shown to be important for tRNA binding (9–11). Specifically, C56 of T-box stacks on a nucleobase in the D-loop of tRNA, and a point mutation of C56 to U has been shown reduce the tRNA binding affinity by ~40 fold (11). We introduced the same mutation in the T-box_182_ backbone (T-box_C56U_) (Fig. 6A and *SI Appendix*, S1). The smFRET trajectories for tRNA^Gly^ binding to T-box_C56U_ are overall similar to the trajectories for WT T-box_182_, with a majority of traces (73±6%) showing stable binding at 0.7 FRET state, and 13±4% of the traces showing transitions back to the 0.4 FRET state (Figs. 6C and D). T0.7 was estimated to be at least ~23 s (limited by the photobleaching of the fluorophore) (Fig. 6E). Post-synchronized time-evolved histogram on the subset of traces that demonstrated real-time binding shows fast transition to the fully bound state (*SI Appendix*, Fig. S7A). Comparison of tRNA^Gly^ binding to T-box_C56U_ and T-box_182_ suggest that the C56U mutation does not affect the second binding step. To investigate whether the mutation at the T-loop region affects the first binding step, we analyzed the binding and dissociation of tRNA^ΔNCCA^-Cy5 to T-box_C56U_-Cy3(3’). We found that the *k*_1_ of tRNA binding to T-box_C56U_ was roughly 16-fold slower compared to tRNA binding to T-box_182_, and the dissociation was roughly 2.5-fold faster compared to WT T-box (Fig. 6E and *SI Appendix*, Fig. S7), leading to a ~40 fold higher dissociation constant for the first binding step. Our results suggest that the T-loop region of the T-box is critical during the first binding step, potentially aiding in anticodon recognition, but does not contribute significantly to the second binding step.

**Figure 6.**
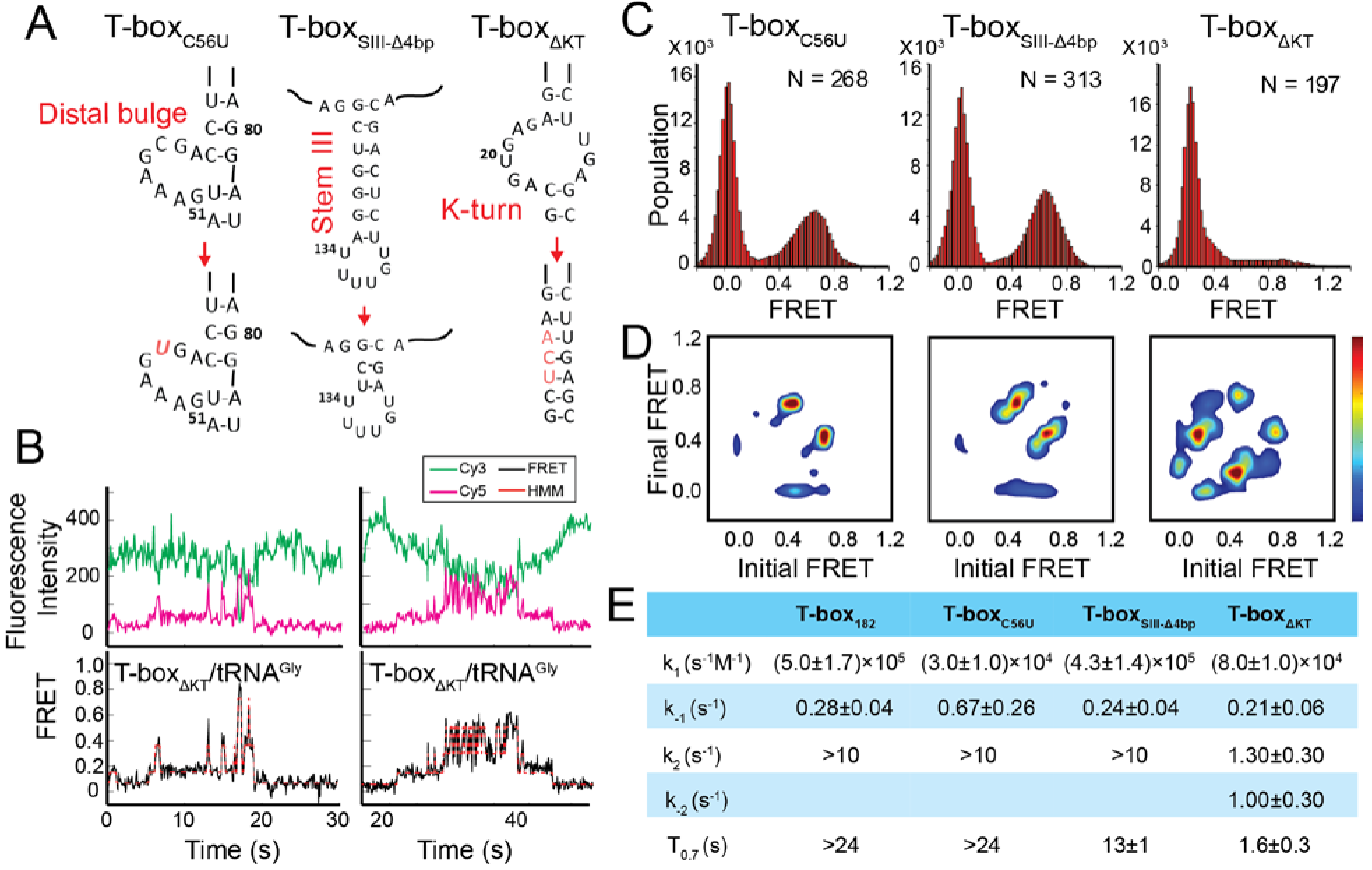
Regulation of the tRNA^Gly^ binding kinetics by structural elements in the *glyQS* T-box riboswitch. **(A)** Schematic diagram of three different mutations introduced to the T-box_182_ backbone (T-box_C56U_, T-box_SIII-Δ4bp_ and T-box_ΔKT_). **(B)** Representative smFRET traces of T-box_ΔKT_-Cy3(3’) and tRNA^Gly^-Cy5. **(C)** FRET histograms of the T-box mutants with tRNA^Gly^-Cy5. **(D)** TDP of the T-box mutants with tRNA^Gly^-Cy5. (E) Table of kinetic parameters for tRNA^Gly^-Cy5 binding to different T-box constructs. *k*_−1_, *k*_2_, and *k*_−2_ of T-box_ΔKT_-Cy3(3’) are apparent rate constants estimated to allow comparison as described in *SI Appendix*.

### A truncation of Stem III has a minor effect on tRNA binding

The functional role of Stem III is unclear. It has been speculated that Stem III might serve as a transcription stalling site to allow co-transcriptional folding and regulation of the T-box riboswitch (21, 22). In addition, a SAXS data-derived model suggested coaxial stacking of Stem III and the anti-terminator stem, leading to a plausible role of Stem III in stabilizing the anti-terminator conformation in the presence of uncharged tRNA^Gly^ (17). To investigate the latter hypothesis, we generated a T-box mutant (T-box_SIII-Δ4bp_), in which four base pairs in Stem III are deleted to significantly shorten its length. smFRET studies using T-T-box_SIII-Δ4bp_-Cy3(3’) with tRNA^ΔNCCA^-Cy5 and tRNA^Gly^-Cy5 revealed insignificant difference in overall kinetics in the first and second step bindings (Fig. 6 and *SI Appendix*, Fig. S8A). Noticeably, the T0.7 was around 50% shorter than that for the T-box_182_ (Fig. 6E), indicating that Stem III may contribute to the stabilization of the fully bound state, potentially through coaxial stacking with the anti-terminator stem, but the effect is minor.

### A K-turn mutation affects both binding steps

As our smFRET data highlight the role of a region near the K-turn as the hinge of the tRNA binding-dependent conformational change, we investigated the role of the K-turn in regulating tRNA binding kinetics. We disrupted the K-turn (T-box_ΔKT_) by changing the 6 bulged nucleotides to 3 nucleotides (UCA) to replace the K-turn with a 3 base pair stem (Fig. 6A and *SI Appendix*, S1). In contrast to binding of tRNA^Gly^ to T-box_182_, binding to T-box_ΔKT_ binding results in three FRET states centered on 0.2, 0.4, and 0.7. (Fig. 6B). While the exact boundary of each FRET state is difficult to determine accurately from the FRET histogram (Fig. 6C), a transition density plot (TDP) clearly revealed interconversion between the 0.2, 0.4, and 0.7 states (Fig. 6D), with transitions between the 0.2 and 0.4 FRET states, and between the 0.4 and 0.7 FRET states more populated. Binding of tRNA^ΔNCCA^-Cy5 to T-box_ΔKT_, on the other hand, leads to the loss of population of the 0.7 state; however, both the 0.2 and 0.4 FRET states and fluctuations between these two states are frequently sampled (Fig. S9).

Comparing the tRNA^Gly^ and tRNA^ΔNCCA^ binding, we speculate that both the 0.2 and 0.4 FRET states observed in the case of the T-box_ΔKT_ represent the partially bound state in which only the anticodon and elbow are recognized. In contrast to T-box_182_, T-box_ΔKT_, with the K-turn replaced by an extension of Stem I, could potentially favor a relaxed conformation of Stem I, as observed in the NMR structure of an isolated K-turn and specifier loop domain (23), generating a lower FRET value centered at 0.2. However, the sampling of the 0.4 FRET in the T-box_ΔKT_ construct suggests that the interactions between the specifier and anticodon of the tRNA may transiently force open the extended base pair region and bend the T-box to adopt a similar conformation to the one observed in T-box_182_. The lifetimes of the 0.4 state transition back to the 0.2 state and the transition forward to the 0.7 state are 0.30±0.03 s and 0.13±0.05 s, respectively, supporting the hypothesis that this forced bent state is energetically unfavorable. However, this 0.4 FRET state is very likely to be required for the NCCA/t-box interaction to occur, as in the presence of tRNA^Gly^ the transitions from the 0.2 FRET to the 0.7 FRET state often pass through the 0.4 FRET state (Fig. 6B). Furthermore, we observed a small region in the TDP corresponding to direct transitions between the 0.2 and 0.7 states. Given the very short lifetime of the 0.4 state, which is close to the time resolution of our experiments, the 0.2 to 0.7 state transition is likely to represent populations whose 0.4 FRET state lifetime is even shorter than the time resolution of the experiment.

The 3-step kinetic scheme for tRNA binding to T-box_ΔKT_ is presented in *SI Appendix*, Fig. S9C. The association rate constant *k*_1_ ((8.0±1.0)10^4^ s^−1^·M^−1^), estimated from binding of tRNA^ΔNCCA^ to the T-box_ΔKT_, is ~ 6-fold smaller than binding to T-box_182_, suggesting that disruption of the K-turn affects anticodon recognition. Considering both the 0.2 and 0.4 FRET state as the partially bound state in T-box_ΔKT_, the apparent dissociation rate from the partially bound state *k*_−1_app_ of tRNA^ΔNCCA^ to T-box_ΔKT_ is 0.21±0.06 s^−1^, similar to that for T-box_182_, suggesting that disruption of the K-turn does not affect the stability of the partially bound state. The 0.7 FRET state observed for T-box_ΔKT_ in the presence of tRNA^Gly^ is consistent with the FRET value for the fully bound state in the WT T-box, indicating that the NCCA/t-box interactions in T-box_ΔKT_ can still be formed. However, tRNA^Gly^ bound to T-boxAKT (T0.7 = 1.6±0.3 s) is at least 10-fold less stable compared to tRNA^Gly^ bound to T-box_182_ (T_0.7_ >24 s). Furthermore, In contrast to the signal observed for tRNA^Gly^ binding to T-box_182_, in which fewer than 10% of the FRET traces show transitions back to 0.4 FRET, in T-box_ΔKT_ the vast majority of the traces show backward transitions to the 0.4 and 0.2 FRET states, contributing largely to the instability of the fully bound state. Based on the transition rates between 0.2, 0.4, and 0.7, we estimated the apparent forward (*k*_2_app_) and (*k*_2_app_) reverse transition rates between the partially bound and the fully bound state of the T-box_Δ_KT to be 1.3±0.3 s^−1^ and 1.0±0.3 s^−1^, respectively (Fig. 6E and *SI Appendix*, Fig. S9C). The dramatically reduced *k*_2_app_ (~100-fold slower than *k*_2_ of T-box_182_) and increased *k*_2_app_ in T-box_ΔKT_ implies that inflexibility of the K-turn region largely inhibits the conformational change in the T-box required to form the NCCA/t-box interaction, and strongly destabilizes the fully bound state.

## DISCUSSION

tRNA recognition by a T-box riboswitch is a bipartite process. Stem I is largely responsible for discriminating non-cognate tRNAs, while the t-box sequence in the expression platform senses the charged state of the tRNA. Here, we used smFRET to elucidate the binding kinetics of tRNA^Gly^ by the *glyQS* T-box riboswitch. With three FRET pairs between different T-box riboswitches and tRNA ligands, our data collectively reveals a two-step binding model of uncharged tRNA^Gly^ to the *glyQS* T-box riboswitch (Fig. 5). The first binding step involves recognition of the anticodon of the tRNA by the specifier sequence located in Stem I of the T-box riboswitch, leading to a partially bound state. In the second step, the 3’ end of the T-box docks into the NCCA end of the tRNA through interactions with the t-box sequence, which leads to a fully bound state. Without the NCCA interaction, the binding of tRNA is unstable, with an average lifetime of ~4 s, whereas with interactions both with the anticodon and the NCCA end, the binding of tRNA is very stable, with an average lifetime > 24 s. The later lifetime measurement is limited by fluorophore photobleaching and is likely to be much longer.

While our manuscript was in preparation, Suddala *et al.* (24) reported a single-molecule study on tRNA binding to the *glyQS* T-box riboswitch and proposed a two-step binding model, very similar to our model. Although both studies propose highly consistent kinetic models, in the study by Suddala *et al.* (24), the FRET pair attached at the variable loop of the tRNA and the 3’ or 5’ ends of the *glyQS* T-box cannot distinguish between the partially bound state from the fully bound state; therefore the two binding states are distinguished by different dissociation rates of the tRNA from these states, aided by using a Stem I-only mutant that cannot interact with the NCCA end of the tRNA. Neither transitions between the partially bound and the fully bound states, nor the order of events during tRNA binding can be resolved in the study (24). In contrast, by employing a FRET pair located at the 5’ end of the tRNA and the 3’ end of the *glyQS* T-box, we observed directly two FRET states corresponding to the recognition of the anticodon (0.4 FRET) and the binding of the NCCA (0.7 FRET), therefore our study allows the discrimination between different states and generates a more complete kinetic framework describing the full trajectory of the tRNA binding process. Specifically, our data reveal that anticodon recognition precedes the NCCA end interactions, and that after anticodon recognition the commitment to further establishment of the NCCA/t-box interaction is high. The T-box/tRNA^Gly^ complex transits rapidly from the partially bound state to the fully bound state, with a rate constant (*k*_2_) of 10 s^−1^. Interestingly, our data also reveal that in the fully bound state the NCCA/t-box interaction is not highly-stable or ultra-stable. Brief disruption of the NCCA/t-box interactions can occur, but transition back to the fully bound state is rapid, ~10 fold faster than dissociation of the tRNA from the partially bound state; therefore tRNA can remain bound during the breaking and reforming of the NCCA/t-box interaction. However, overall, such transient breaking of NCCA/t-box interaction was only observed in ~10% of the total population, suggesting that the reverse transition rate constant (*k*_−2_) is very small, and the second binding step in the WT T-box with uncharged tRNA^Gly^ is close to be irreversible.

The first binding step, interactions with Stem I, involves more than anticodon recognition. Crystal structures of Stem I in complex with tRNA (11, 14) show the predicted interaction between the interdigitated T-loops in Stem I and the elbow region (D- and T-loops) of the tRNA. Our T-box/tRNA FRET pairs and the intra-T-box FRET pair do not report directly on the formation of T-loops/elbow interactions, and therefore cannot resolve whether the T-loops/elbow interactions proceed before or after anticodon recognition. Nevertheless, the importance of this interaction is captured by smFRET measurements using a point mutation that impairs this interaction (11). The mutant shows a dramatic decrease in the association rate constant and a moderate increase in the dissociation rate constant, leading to an overall ~40 fold reduction on the binding affinity for the first step, but the second step is unaffected, suggesting that establishment of the T-loops/elbow interactions is an important part of the Stem I/tRNA recognition process, but plays a minimal role in NCAA recognition or binding.

Consistent with the study by Suddala *et al.* (24), a FRET pair at the 5’ and 3’ ends of the T-box generates a high FRET value that is insensitive to any tRNA binding, suggesting that the T-box is largely pre-organized in a folded state before tRNA binding. However, by adding an additional extension sequence at the 3’ end, we find that the 3’ half of the T-box (including Stem III and the anti-terminator) moves inward relative to the 5’ half (Stem I) of the T-box to accommodate the interaction with the NCCA end. The observation that such conformational change is only associated with the presence of tRNA^Gly^ indicates that this intra-T-box conformational change is a concerted motion to aid in the establishment of the NCCA/tbox interaction. From the FRET pair placed on the tRNA and at the 5’ end of Stem I, we confirm that the hinge is likely to be near the K-turn region. Furthermore, the crystal structures (11, 14) show that Stem I flexes around the K-turn region and this flexing seems to be important to establish the interactions between the anticodon and specifier sequence. Our smFRET experiments using a mutant where the K-turn region is removed show that in the absence of the K-turn motif, both the first and second binding steps are significantly affected, emphasizing the importance of the flexing around the K-turn region in the overall recognition and binding process. Interestingly, Stem III does not appear to play a major role in the tRNA binding process *in vitro*, with a minor effect on the stability of the fully bound state. Stem III may contribute to the stabilization the anti-terminator conformation *in vitro*; however, potentially its major function is to create a pause site to coordinate with the co-transcriptional folding of the T-box (21, 22).

## MATERIALS AND METHODS

T-box and tRNA samples (*SI Appendix*, Fig. S1) were prepared by *in vitro* transcription and purified by 8M urea polyacrylamide gel electrophoresis (PAGE) ((17) and *SI Methods*). End labeling of the T-box constructs and the tRNAs was performed based on published protocols with modifications ((25) and *SI Methods).* smFRET experiments were performed in 20 mM HEPES pH 7.0, 100 mM KCl and 15 mM MgCl_2_ supplemented with 5 mM protocatechuic acid (PCA) (Sigma), 160 nM protocatechuate-3,4-dioxygenase (PCD) (Sigma) and 2 mM Trolox (Sigma). Images were recorded with an objective-based total internal reflection fluorescence (TIRF) microscope with 100X NA 1.49 CFI HP TIRF objective (Nikon). A 561 nm laser (Coherent Obis at a power density of 4.07 × 10^5^ W/cm^2^) was used for the FRET measurement. Individual spots were picked from maximum projection of Cy5 emission channel by NIS Elements software. Cy3 and Cy5 fluorescence intensity *vs.* time trajectories were corrected for baseline and bleed-through in MATLAB as previously described (26) and fit with a Hidden Markov Model using vbFRET (27). Dwell times of each FRET state before transition to another FRET state were extracted from the idealized traces and fit with a single or double exponential decay with Origin 7.0 (OriginLab). Full methods and references can be found in the *SI Appendix*.

## ACKNOWLEDGEMENTS

This work was funded by a grant to JF and AM by the Chicago Biomedical Consortium with support from the Searle Funds at the Chicago Community Trust. We thank E. Heideman for the preparation of slides for all imaging experiments and managing the Fei laboratory and Dulce Alonso Hernandez for help and advice with RNA labeling.

**AUTHOR CONTRIBUTIONS:** JZ, BC, AM, and JF designed the research. JZ conducted smFRET experiments and analyzed the data. BC prepared the samples and conducted ITC experiments. EC helped with imaging and data analysis. WL and JF helped with data analysis. JZ, BC, AM, and JF wrote the manuscript; all authors approved the final manuscript.
The authors declare no conflict of interest.

